# SXTractor: A Self-Supervised Feature Extractor of Soft X-Ray Images That Enables Few-Shot Tomogram Segmentation

**DOI:** 10.1101/2025.08.25.672073

**Authors:** ShaoSen Chueh, Madeleen C. Brink, Jeremy C. Simpson, Sergey Kapishnikov

## Abstract

Soft X-ray tomography (SXT) is a powerful, non-invasive bio-imaging technique that enables visualization of cellular structures in near-native states. Despite its potential, the development of dedicated image analysis tools — particularly deep-learning-based models — has been limited, largely due to the limited accessibility of soft X-ray microscopes and the scarcity of labeled SXT data. To address this deficit, in this work, we present SXTractor, a self-supervised SXT feature extractor based on the DINO framework. SXTractor can be fine-tuned with minimal labeled data and effectively adapted to various downstream tasks. We demonstrate its utility on few-shot tomogram segmentation, where it significantly outperforms the model when trained from scratch. Furthermore, it achieves few-shot segmentation performance comparable to that of the Segment Anything Model (SAM), despite SAM being a segmentation-specific model pretrained on millions of labeled images with a significantly larger model size. Most importantly, SXTractor enables a diverse range of downstream applications of deep learning to SXT, thus offering a practical and scalable solution for SXT image analysis in data-constrained settings.

## Introduction

Soft X-ray tomography (SXT) is a powerful bio-imaging technique that provides unique advantages for imaging near-native cryo-biological specimens. Its resolution capabilities effectively bridge the gap between light and electron microscopy, making it a valuable complementary tool to these established imaging modalities [1][2]. Despite its advantages, the widespread adoption of soft X-ray tomography is hampered by limited access to the beamline sources. The majority of soft X-ray sources have previously been confined only to stadium-sized synchrotrons, which significantly restricts their availability for researchers. In recent years, laboratory-based soft X-ray microscope systems have been developed [3][4][5], which are expected to unlock the potential of this transformative imaging technique.

However, currently, the limited amount of Soft X-ray Tomography (SXT) data available has led to a significant lack of dedicated image analysis tools. This issue has become even more pronounced with the rapid advancements in deep-learning-based bioimage analysis methods [6], particularly for segmentation in microscopy images [7] [8]. A major hurdle for deep learning segmentation models is their reliance on large amounts of data labeled by human experts. Such datasets are rarely available to most researchers working with SXT. To make matters worse, the diversity of cell types and imaging conditions makes it challenging for these deep learning models to generalize effectively across different datasets. For instance, MitoNet [9], a mitochondrial segmentation model for electron microscopy, required over a million 2D image patches for training. This scale of data is not currently feasible for SXT, highlighting a critical need for new approaches to image analysis in this field.

The Segment Anything Model (SAM) [10] presents a novel paradigm for developing segmentation models. SAM is a foundation model trained on an unprecedented 11 million images (the SA-1B dataset) with a total of approximately 93 million parameters, providing a robust base for fine-tuning across downstream segmentation tasks. Building upon this, models such as Segment Anything for Microscopy [11] and Segment Anything in Medical Images [12] have demonstrated the adaptability of the SAM architecture to specialized domains. However, SAM’s architectural design, which incorporates prompting encoders in addition to the standard image encoder and decoder layers, may render it less than optimally suited for direct fine-tuning towards fully automatic segmentation. Furthermore, SAM’s domain specificity to segmentation poses a challenge for wider applications; developing similarly comprehensive foundation models for every distinct image processing task may not be the optimal solution for microscopy image processing.

On the other hand, self-supervised learning has emerged as a profoundly insightful approach within the field of computer vision [13][14][15][16]. Notably, the self-supervised framework DINO (Self-***Di***stillation with ***No*** Labels) [17][18] has extensively explored self-supervised training techniques for vision transformers (ViTs). This research has revealed a remarkable property: a self-supervised Vision Transformer (ViT) is capable of learning the semantics of images without any explicit labels. This discovery of inherent semantic understanding, cultivated through self-supervised pretraining, offers a compelling new perspective on the application of vision transformer models for various downstream tasks.

Given the novel methodologies introduced and the present scarcity of SXT data, few-shot tomogram segmentation offers a more generalizable and robust solution for SXT image analysis than developing highly specialized organelle-specific models, which often struggle to generalize across datasets. This approach allows biologists to reliably label only a few representative slices (5∼10 slices in our experiments) with their expertise knowledge of the specimens, while deep learning models, fine-tuned with the limited number of labeled slices, can efficiently segment the remaining slices, thereby significantly reducing manual annotation effort. Prior research has robustly demonstrated the applicability of few-shot learning to a variety of image segmentation tasks [19][20][21][22][23]. Critically, most successful implementations of few-shot segmentation leverage techniques such as self-supervised learning, meta-learning, or foundation models like SAM, enabling models to rapidly adapt to novel tasks with only a limited amount of annotated data. This alignment between existing few-shot strategies and the characteristics of SXT data strongly supports its adoption in this domain.

In this study, we introduce SXTractor, a DINO-based self-supervised pretrained SXT feature extractor. SXTractor aims to establish a robust foundation for subsequent fine-tuning across a diverse array of SXT image processing tasks, including, but not limited to, few-shot tomogram segmentation. Our findings demonstrate that leveraging this pre-trained feature extractor significantly reduces the amount of labeled data required for fine-tuning, leading to substantial performance improvements compared to the model (with the same architecture) trained from scratch. Our study further demonstrates SXTractor’s efficacy by comparing its few-shot tomogram segmentation performance with that of a fine-tuned Segment Anything Model (SAM). Remarkably, SXTractor achieves the same level of accuracy in few-shot segmentation settings. This parity in performance highlights SXTractor’s significant potential: it can accomplish downstream few-shot segmentation tasks comparable to those of a gigantic foundation model (such as SAM with ∼93 million parameters trained on 11 million 2D labeled images), yet with a substantially smaller pretraining dataset (less than 5,000 2D unlabeled tomogram slices) and number of parameters (∼21 million). Furthermore, different from SAM, which is a segmentation-specific foundation model, SXTractor is a non-task-specific encoder, making it highly adaptable to a wide range of image processing tasks beyond just segmentation for SXT.

## Results and Discussions

### SXTractor: DINO-Based Pretraining of SXT Feature Extractor

The DINO framework leverages both self-supervised learning and knowledge distillation [24][25][26]. Instead of using a pretrained model as the teacher network, DINO uses a dynamically updated teacher network with an exponential moving average (EMA) from the student parameters (Fig. 1a). After 2 different augmentation strategies (global and local augmentations, for passing into the networks), DINO aims to train the student network to match the output of the teacher network, forcing it to learn the core image features. DINO has shown that this method can extract core image semantics, which has not been seen in previous supervised-trained vision transformer models [27] or convolutional neural networks (CNNs). Our results (visualization of attention heads) also verify that SXTractor, based on the DINO framework, can extract cellular features of SXT images without being given any labels for supervised training (Fig. 1b). Interestingly, the visualization also reveals that SXTractor exhibits a tendency to recognize and de-emphasize the missing wedge artifact [28] in SXT images, placing reduced attention on the elongated regions affected by this distortion. This characteristic proves critical for accurate few-shot segmentation: the SXTractor-based model generates substantially fewer false positives compared to the model trained from scratch in the area with missing wedge artifacts. This improvement can be attributed to SXTractor’s capacity to distinguish blurred or distorted structures caused by the missing wedge and to avoid erroneously labeling these ambiguous regions.

**Figure 1.**
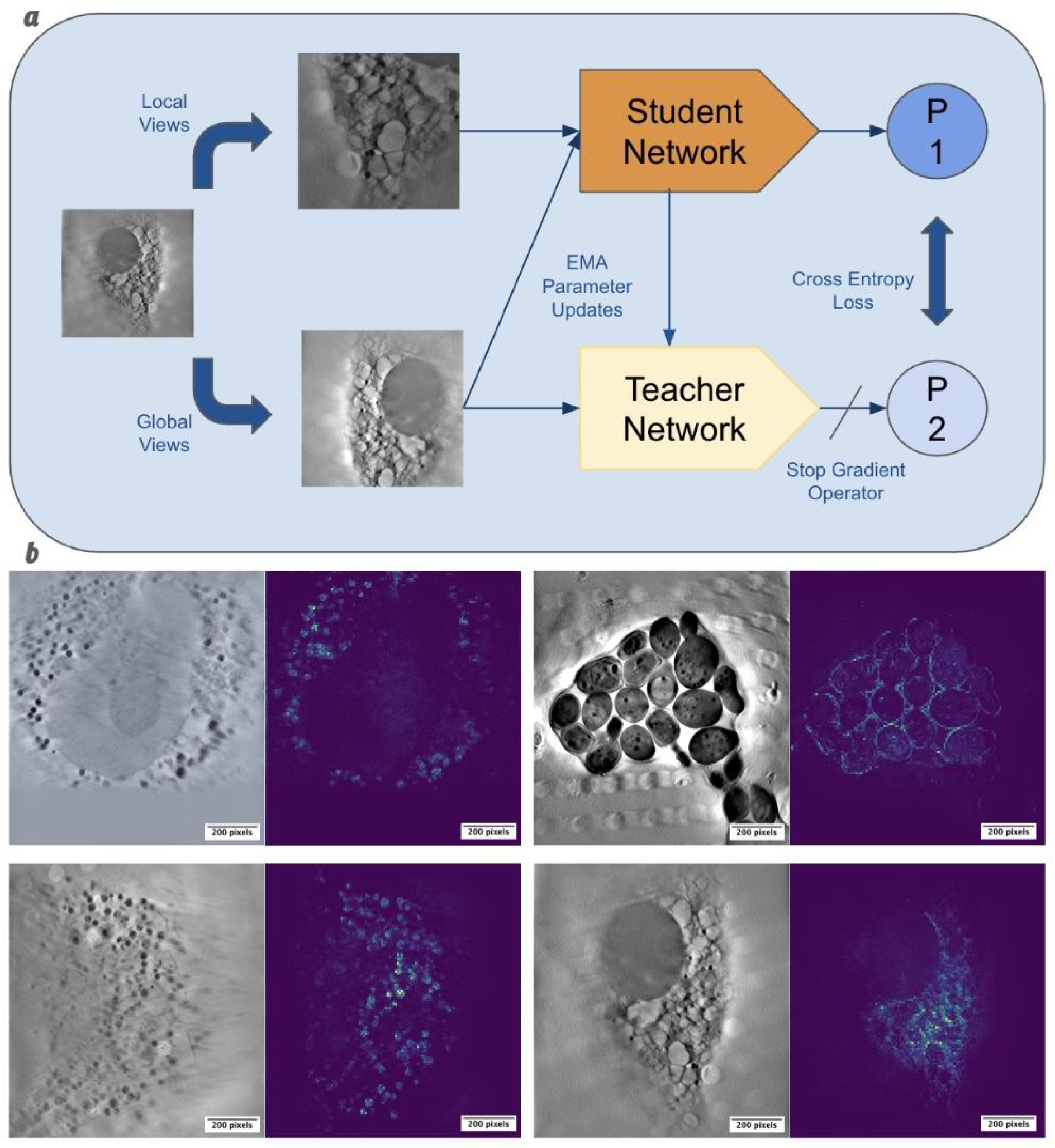
SXTractor pretraining based on the DINO framework (Pixel Size: 28.8 nm). The brighter pixels (higher pixel values) indicate a higher attention output by the model. **a)** The DINO framework applies 2 different augmentation strategies (global and local) to an SXT image that contains cellular structures, then puts them into the student and teacher networks (only global views go into the teacher network, while both views are passed into the student network). By forcing the output of the student network to match the output of the teacher network, the model learns to extract the core features of SXT images. **b)** The visualization of attention maps shows SXTractor’s capability of extracting key semantic features of diverse cellular structures in SXT images. (Top-left: VeroE6, Top-right: Yeast cells BY4741, Bottom: HeLa cell)

We are also curious about how useful transfer learning [29] is for a dataset in a different domain. We trained the SXTractor both with pretrained DINO parameters and from scratch to observe the difference in performance. The result shows that training with pretrained DINO parameters converges much faster and reaches a lower level of loss (constantly below 1.0 within 150 epochs), while training from scratch converges much slower (about 1500 to 2000 epochs), and the loss fails to decrease below 1.0. We visualize the attention layers of both training schemes in Fig. 2a. It can be seen that the output of the model trained from scratch (Fig. 2a, right column) still demonstrates the capability of putting higher values (attention) on the pixels of key cellular structures. However, the background noise level is much higher compared to the model trained with pretrained DINO parameters (Fig.2a, mid column).

**Figure 2.**
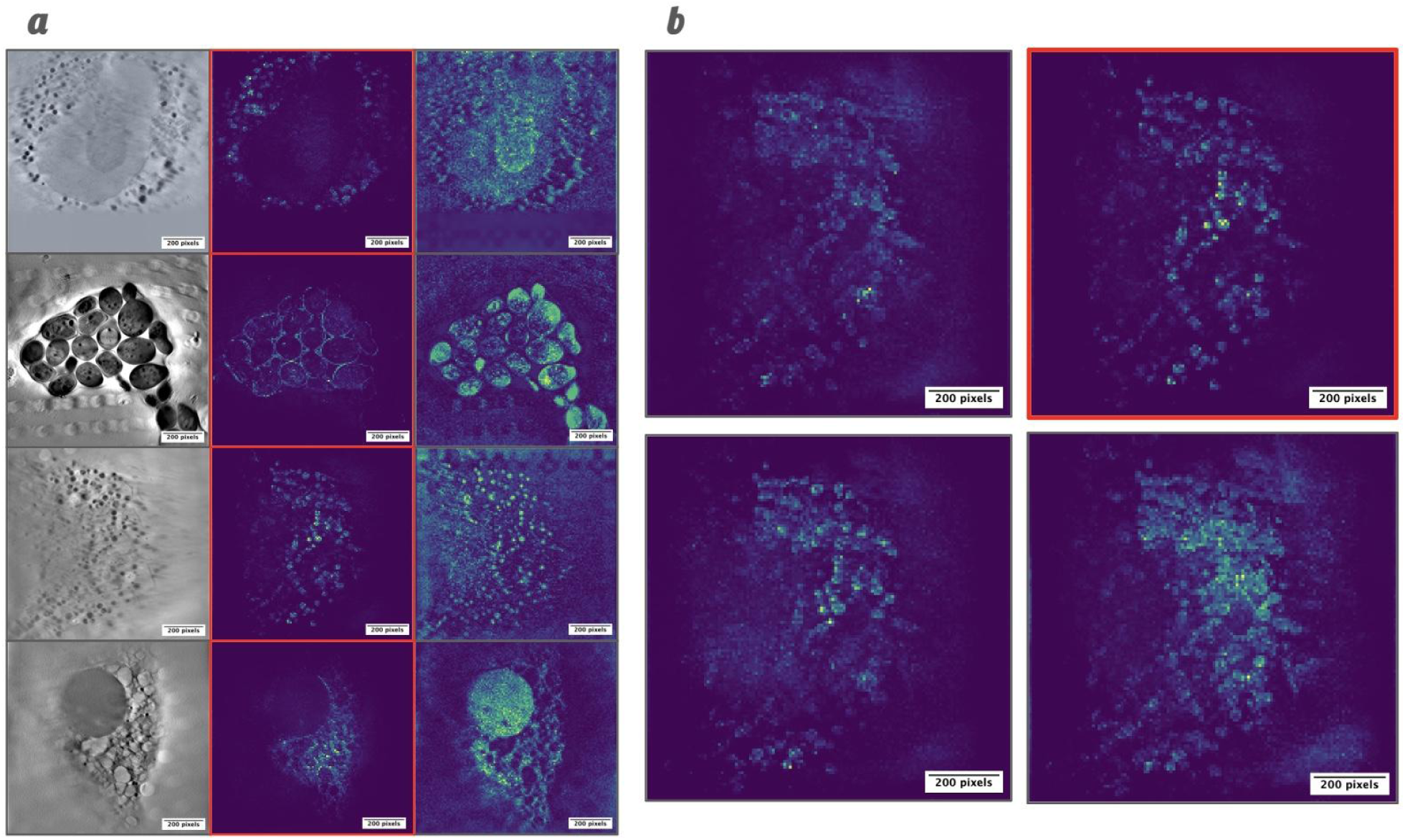
Comparison of SXTractor with different training strategies (Pixel Size: 28.8 nm). **a)** Attention maps of the models trained with pretrained DINO and from scratch(Cells from top to bottom: VeroE6, Yeast cells BY4741, and HeLa Kyoto.). This visualization shows that the model based on pretrained DINO parameters is able to minimize the background noise. (Left: SXT image, Mid: trained with pretrained DINO, Right: trained from scratch). **b)** Attention maps across training epochs reveal that longer training does not necessarily yield better results (HeLa Cell). This may be due to overfitting on the limited SXT dataset, which gradually diminishes the benefits of the pretrained DINO parameters.

Lastly, the SXTractor model was trained for 400 epochs with pretrained DINO parameters. While the training loss decreased below 1.0 consistently after 100 epochs, a qualitative assessment of the visualized attention maps indicated that a longer training duration does not necessarily correspond to improved visual clarity. Visualizations of the model’s performance at epochs 100, 200, 300, and 400 (Figure 2b, upper left, upper right, lower left, and lower right, respectively) revealed that the cellular structures were most distinctly delineated from the background at the 200th epoch. Conversely, extending training to 300 and 400 epochs appeared to introduce increased background noise. This phenomenon suggests that continued training on limited SXT data may lead to overfitting and that the representational power of the pre-trained DINO parameters may be diminishing.

Consequently, the model checkpoint saved at the 200th epoch was selected as the final SXTractor model due to its optimal performance in terms of visual quality.

### SXTractor for Few-Shot Soft X-Ray Tomogram Segmentation

#### 1. Challenges in SXT Segmentation

Segmentation of cellular structures in soft X-ray tomography (SXT) is inherently challenging due to several factors. First, as a newly emerging imaging modality, SXT lacks the extensive publicly available datasets that exist in those more established techniques, such as electron or light microscopy. This scarcity of annotated data limits the effectiveness of supervised deep learning approaches. Second, the presence of the missing wedge artifact in tomographic reconstructions complicates the labeling process[28]. As illustrated in Fig. 4a, the resulting elongation and distortion of cellular structures make it difficult to distinguish objects, even for expert biologists and imaging physicists. This, in turn, hinders model training and contributes to lower segmentation accuracy, as demonstrated in our experiments. Lastly, the high variability in the morphology of cells and organelles — where the same organelle may appear markedly different across cell types and states — poses a significant challenge for generalization. To address these issues, we adopt a few-shot learning strategy rather than developing a collection of organelle-specific segmentation models.

**Figure 3.**
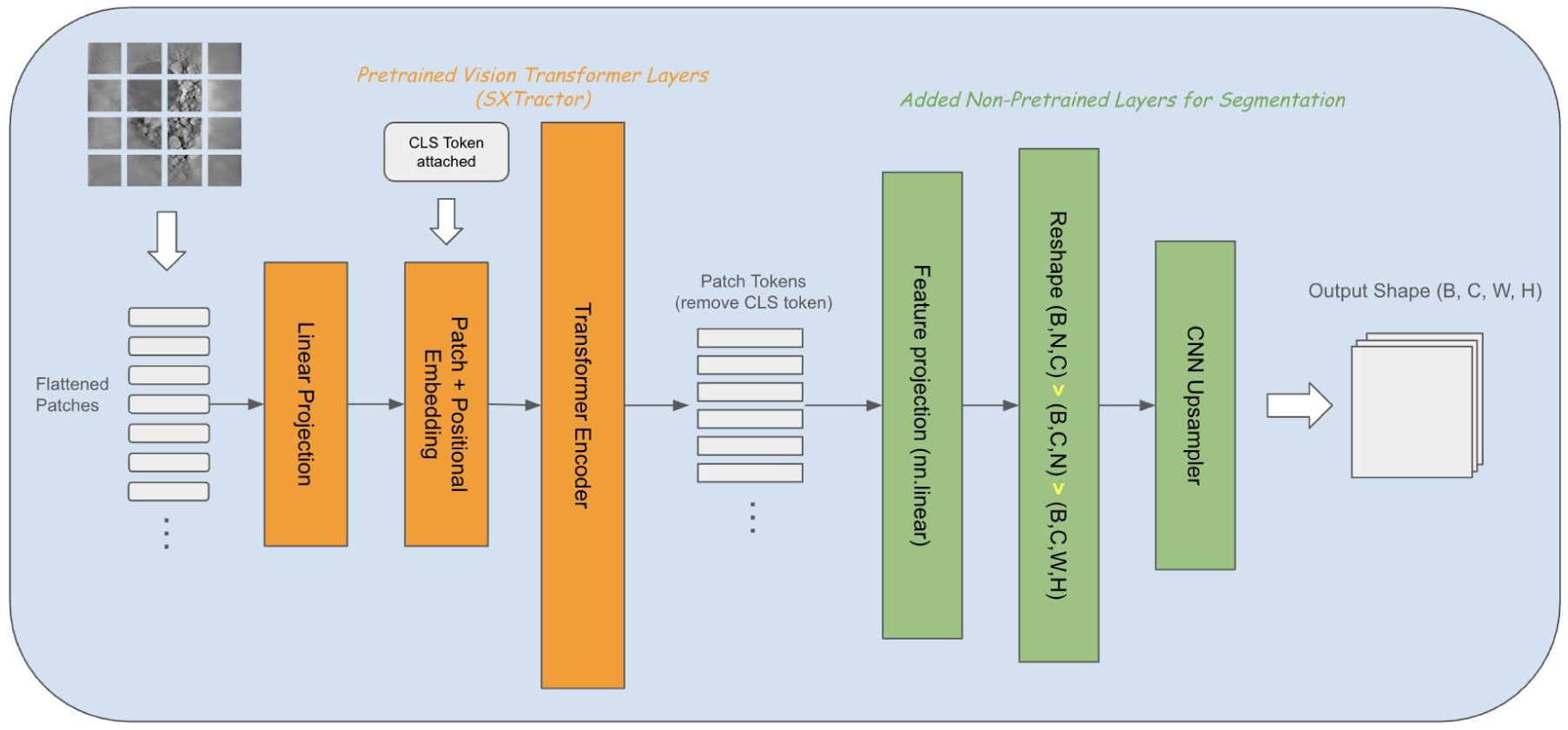
Segmentation model architecture (SXTractor + Upsampling CNN Layers). The architecture is based on the pretrained encoder SXTractor with the convolutional decoder for segmentation. The output of SXTractor is of dimension (B, N, C) instead of the conventional 2D tensor as in common vision models. After reshaping the dimensions from (B, N, C) into (B, C, W, H), the convolutional upsampling layers are applied to upsample the dimensions back to the original input image shape.

**Figure 4.**
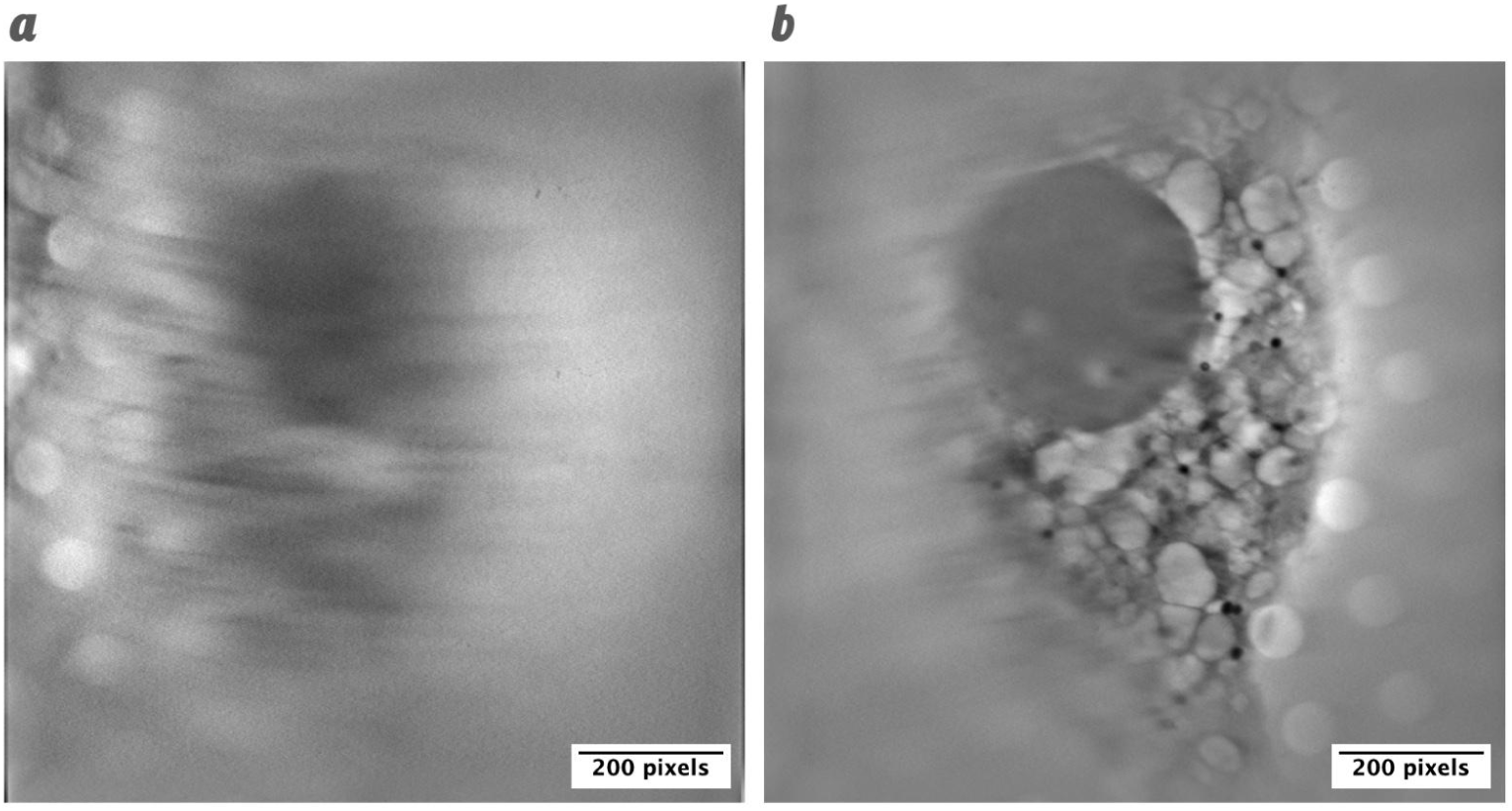
Distortions caused by the missing wedge (a) in HeLa Cell (Pixel Size: 28.8 nm). The distortions caused by the missing wedge effect make consistent labelling of SXT data challenging.

#### 2. Few-Shot Segmentation with SXTractor

Previous studies have demonstrated the versatility of DINO-based models across a range of visual tasks [30][31][32]. Building on this, we utilize our SXTractor model, along with a CNN-based decoder, for few-shot tomogram segmentation. Following DINO-based pretraining (Fig. 1), SXTractor learns to extract informative cellular features from soft X-ray tomography (SXT) images, thereby establishing a strong foundation for downstream segmentation tasks. However, unlike conventional convolutional neural networks (CNNs) [33][34][35][36][37], which produce spatially-structured feature maps with shape (Batch size, Channel, Height, Width), the Vision Transformer architecture outputs a sequence of tokens with shape (B, N, C), where B is the batch size, N is the number of patches, and C is the embedding dimension — functionally analogous to feature channels in CNNs. To interface this output with a CNN-based decoder, it is necessary to first permute the tensor to (B, C, N), and then reshape the sequence dimension N into its original 2D spatial configuration, yielding a tensor of shape (B, C, H, W). This transformation is essential because CNNs require the channel dimension to precede the spatial dimensions and operate on inputs with explicit spatial locality, which is encoded in the sequence ordering of Vision Transformer outputs. The overall segmentation architecture is shown in Fig. 3.

In our experiments, we annotated lysosomes and nuclei in HeLa cells imaged by soft X-ray tomography (SXT), using correlated fluorescence microscopy to improve labeling accuracy. These two organelles were selected because we want to evaluate model performance at different levels of segmentation difficulty. Lysosomes, due to their small size and similarity to other organelles, are particularly challenging to segment, especially under distortion from the missing wedge artifact. In contrast, nuclei are larger and more morphologically distinct, making them easier to identify despite some degree of distortion. Our experimental results also verify this assumption. To evaluate the model under few-shot learning conditions, we trained it using only 5 and 10 annotated slices in a tomogram, while the remaining slices in the tomogram were reserved for evaluation. We note that beyond 10 labeled slices, the practical utility of few-shot learning diminishes, as the remaining slices in the tomogram can often be annotated via interpolation, reducing the need for model-based segmentation.

Despite the use of fluorescence-guided labeling, the missing wedge artifact inherent in grid-based SXT acquisition (Fig. 4) remained a significant challenge. The presence of this artifact, particularly impacting small organelles like lysosomes, complicates accurate annotation and can lead to imperfect ground truth data. Consequently, quantitative metrics such as the Dice score may not fully capture a model’s true performance. To thoroughly assess our proposed model, we conducted both qualitative analysis and a comparative analysis against two baselines: (1) the Segment Anything Model (SAM), a state-of-the-art foundation model for segmentation, and (2) an identical architecture trained from scratch without any pretraining.

Lastly, there has been ongoing discussion in the literature regarding whether freezing some, or all, encoder layers during transfer learning can help mitigate overfitting in low-data regimes [38][39]. However, empirical findings on this topic have been mixed. In our experiments, we observed that freezing the pretrained SXTractor encoder during downstream few-shot segmentation did not yield noticeable improvements in generalization performance. Conversely, allowing the entire model — including both the SXTractor encoder and the CNN-based decoder — to be updated during few-shot training led to a stable decrease in validation loss. Based on these findings, we opted not to freeze any layers of SXTractor during downstream segmentation training.

#### 3. Lysosome Segmentation Results

Our experiment results demonstrate that SXTractor achieves comparable validation accuracy to the Segment Anything Model (SAM) in few-shot lysosomes segmentation scenarios, with both models significantly outperforming the version trained from scratch without pretraining. In the 5-shot experiments (Fig. 5a, validation dice scores), the Segment Anything Model (SAM) demonstrates a faster convergence rate, achieving high validation performance within relatively few training epochs. This behavior is expected, as SAM incorporates both a pretrained encoder and decoder optimized for segmentation tasks (no prompts were used in our experiments). In contrast, our SXTractor-based model employs a general-purpose pretrained encoder (SXTractor) with a randomly initialized decoder, which naturally requires more training to adapt. Nevertheless, after approximately 1000 training epochs, the SXTractor-based model rapidly catches up in terms of validation Dice score, with both models ultimately achieving comparable performance around 60% (Dice score ranges from 0 to 1, and we will refer to it as a percentage of overlap in this study). In comparison, the model trained entirely from scratch without pretraining lags significantly behind. In the 10-shot experiments (Fig. 5b), a similar trend is observed. Both SAM and the SXTractor-based model substantially outperform the model trained from scratch. Notably, the SXTractor-based model achieves a slightly higher validation Dice score than SAM, reaching nearly 70% compared to SAM’s 68%.

**Figure 5.**
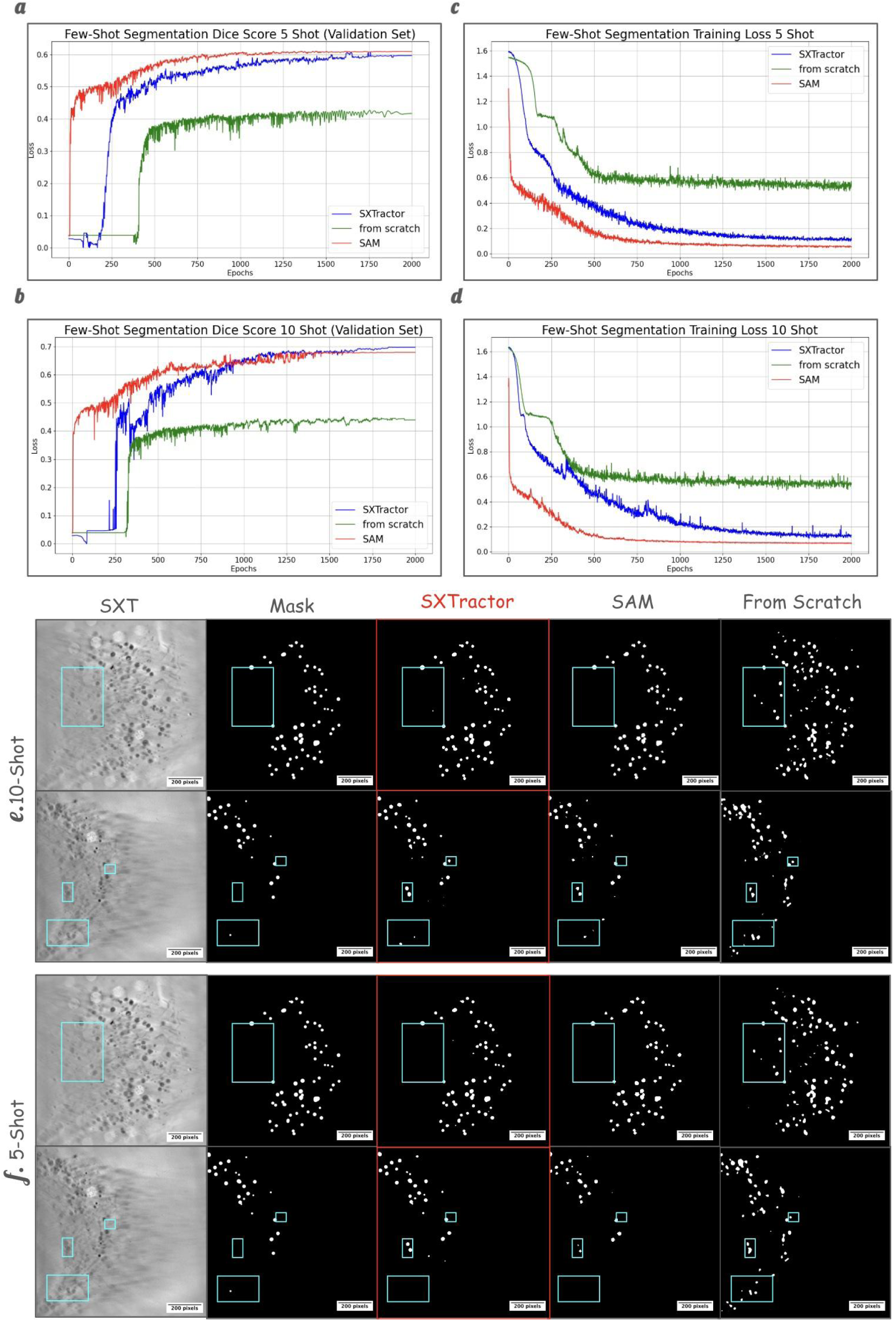
Few-shot lysosome segmentation (Pixel Size: 28.8 nm). **a, b)** Validation Dice Scores. SXTractor-based model and SAM show a significant advantage in segmentation accuracy over the model trained from scratch (same architecture as SXTractor), which struggles to converge within 2000 epochs. **c, d)** The training loss of the trained-from-scratch model shows a slight downward trend, suggesting that its performance could improve with extended training. **e, f)** Qualitative comparison of different models. The predictions of our model and SAM show high similarity to the ground-truth mask, while the model trained from scratch contains more false-positive predictions.

However, it is worth noting that although the model trained from scratch lags in terms of Dice score accuracy after 2000 epochs, its training loss (Fig. 5c, 5d) continues to exhibit a downward trend. This suggests that further training may eventually lead to improved segmentation performance. Nonetheless, training time is a critical consideration in few-shot learning scenarios, as one of the primary motivations for adopting few-shot approaches is to reduce both annotation and computational overhead. Models that require substantially longer training to achieve competitive performance may not be practical in real-world applications where rapid adaptation to new data with minimal supervision is essential.

Although the Dice scores in both the 5-shot and 10-shot experiments do not reach an ideal level above 80–90%, this limitation largely reflects the inherent difficulty of lysosome segmentation in SXT (compared to ∼94% accuracy of nucleus segmentation for both SXTractor and SAM in Fig. 6). Nevertheless, qualitative results (Fig. 5e, 5f) demonstrate strong agreement between the predicted masks (generated by our SXTractor-based model and SAM), and the manually annotated ground truth, albeit with some remaining false positives (Fig. 5e, 5f, boxed area). In contrast, the model trained from scratch — despite sharing the same architecture — exhibits significantly more false positives. Notably, these errors are not randomly distributed; most occur in regions affected by the missing wedge distortion. During annotation, we deliberately avoided labeling structures in these areas due to their ambiguous appearance, particularly for small organelles such as lysosomes and lipid droplets. However, the distortion introduced by the missing wedge is a gradual effect across slices, making it difficult to define a precise threshold for inclusion or exclusion during manual labeling, even for human experts.

**Figure 6.**
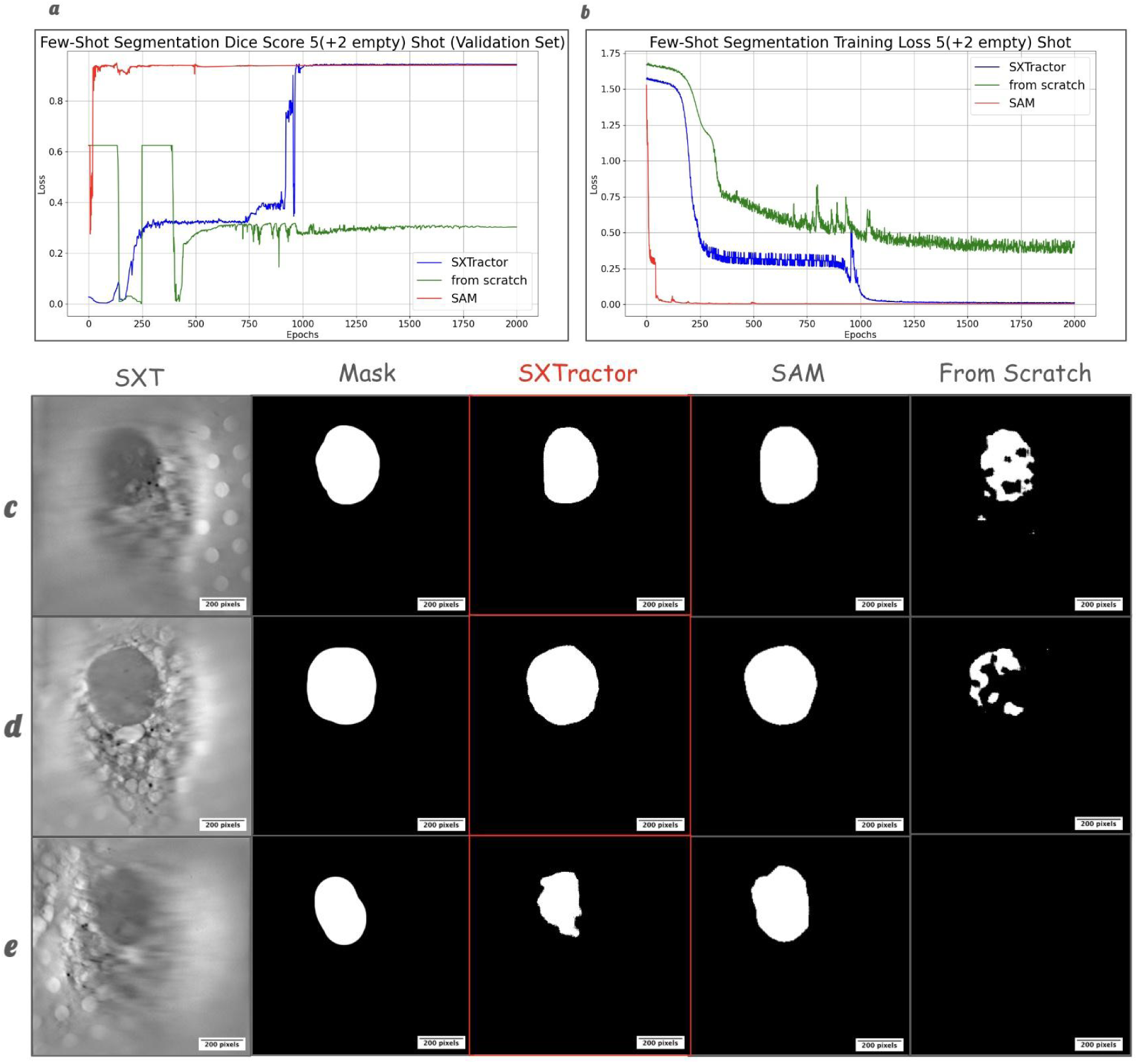
Few-shot nucleus segmentation (Pixel Size: 28.8 nm). a) Validation Dice scores. Both our SXTractor-based model and SAM achieve ∼94% accuracy, while the model trained from scratch fails to converge effectively. b) Training loss. SXTractor-based and SAM models rapidly minimize loss, whereas the model trained from scratch exhibits a much slower decrease in loss, which does not translate into improved Dice score performance. c–e) Visualization of model predictions. In undistorted slices, both our model and SAM produce masks that closely align with the ground truth, while the model trained from scratch displays major inaccuracies. However, all models struggle to make consistent predictions in slices affected by the missing wedge artifact.

As a result, evaluating lysosome segmentation performance using a single metric or strict standard can be misleading. Our annotation strategy focused on labeling only the clearest and most interpretable lysosomes, while disregarding ambiguous, distorted regions. From this perspective, both our SXTractor-based model and SAM more closely align with our labeling goals by avoiding over-segmentation in distorted regions. In contrast, the model trained from scratch tends to over-label these areas, leading to inflated false positive rates and thus lower Dice scores.

#### 4. Nucleus Segmentation Results

Compared to lysosomes, the nucleus is a significantly easier target for segmentation. Although still affected by the missing wedge artifact (Fig. 4), the larger volume and more distinct morphology make them easier to label. Our quantitative results support this assumption: without any data augmentation and within just 1000 training epochs, both the SXTractor-based model and SAM achieve ∼94% Dice score, while the model trained from scratch struggles to converge, plateauing below 35% (Fig. 6a). Although the training loss of the model trained from scratch (Fig. 6b, green line) shows a steady downward trend, this does not translate into improved validation accuracy.

In our experiments, an unusual pattern appears in the Dice score of the model trained from scratch (Fig. 6a), which starts around 60% and then suddenly drops to zero after a few epochs. This initially high score is misleading. Since only ∼40% of the slices in the tomogram contain clear nuclei that we choose to label, the remaining 60% have empty masks. At the beginning of training, the model trained from scratch tends to predict empty outputs for all slices, coincidentally matching the majority of empty ground truth masks and artificially inflating the Dice score. As training progresses and the model begins to make non-empty predictions, the score drops, reflecting the true performance of the model trained from scratch.

Importantly, we observed that the choice of training slices strongly influences performance. Initially, we labeled five slices containing clearly visible nuclei (Fig. 4b), but the validation Dice score stalled around 30-35%, despite training Dice reaching over 90%. Upon inspection, we found the models were predicting nuclei in distorted slices we deliberately left unlabeled. To address this, we added two distorted slices (Fig. 4a) with blank masks to explicitly inform the model of our labeling threshold. With this adjustment — five clear and two blank distorted slices — the validation Dice scores rise above 90% for both pretrained models, showing that a few well-selected training slices can guide model learning effectively.

In our qualitative analysis, we visualize the segmentation outputs of each model in Fig. 6c–e. Both the SXTractor-based model and SAM produce masks that closely match the annotations, while the model trained from scratch shows significant errors. Notably, in slices near the threshold of whether to label — due to missing wedge distortions (Fig. 6e) — both models (ours and SAM) struggle to generate consistent predictions. This is expected, as even expert biologists find it difficult to make definitive labeling decisions in such ambiguous cases. With only a few annotated training slices, the models naturally inherit this uncertainty.

#### 5. Domain Consideration: Grid vs. Capillary

It is worth noting that biological samples for soft X-ray tomography (SXT) are typically prepared using one of two substrate types: electron microscopy (EM) grids or cylindrical capillaries [40][41][42][43]. EM grids are the more established and widely adopted option, also commonly used in other imaging modalities such as electron tomography. However, their planar geometry introduces a significant limitation, namely the missing wedge artifact, which affects the completeness of tomographic reconstructions and complicates downstream segmentation tasks. In contrast, capillary-based imaging circumvents the missing wedge issue due to its cylindrical geometry, resulting in more isotropic data and improved segmentation feasibility [44][45]. Despite this advantage, capillaries are far less commonly used, leading to a severe scarcity of available data. Given these trade-offs, this study focuses on SXT images acquired using EM grids. As a result, the SXTractor model is pretrained and evaluated exclusively on grid-based data. Due to the substantial domain shift between grid and capillary datasets, the current implementation of SXTractor is not expected to generalize effectively to capillary-acquired images.

## Discussion

In summary, rather than developing numerous organelle-specific segmentation models for highly variable microscopy data, a few-shot tomogram segmentation strategy offers a more efficient and generalizable solution. Although organelle-specific models could fully automate the segmentation, the limited volume of SXT data makes it unrealistic to expect strong generalization across diverse datasets. On the other hand, although few-shot tomogram segmentation requires users to label a few slices, it significantly improves the reliability of the model predictions because the variability is much smaller within the same tomogram (and the same cell). Furthermore, a few-shot tomogram segmentation model can be applied to organelles of different sizes and shapes (as in our experiments: lysosome and nucleus), eliminating the need to develop separate models for each organelle and substantially reducing development effort.

Our results highlight the importance of robust pretrained models for effective segmentation in few-shot learning scenarios (Fig. 7). In domains such as soft X-ray tomography (SXT), where annotated data are scarce and labeling demands substantial domain expertise, a strong pretrained backbone greatly reduces the need for extensive manual annotation. While both SAM and SXTractor achieve comparable segmentation accuracy, their characteristics differ markedly. SAM, a large segmentation-specific foundation model (∼93 million parameters, based on the smaller version ViT-B) trained on millions of annotated natural images, benefits from task specialization and converges quickly but demands substantial GPU memory and computational resources. In contrast, the SXTractor-based segmentation model — a compact general-purpose encoder (∼21.8 million parameters) pretrained with self-supervised learning on fewer than 5,000 unlabeled SXT slices — offers a lighter-weight, resource-efficient alternative. Its smaller size makes it practical for laboratories with limited computational capacity, and most importantly, its general-purpose design enables potential application to a wide range of SXT image analysis tasks beyond segmentation, making it a scalable and versatile solution.

**Figure 7.**
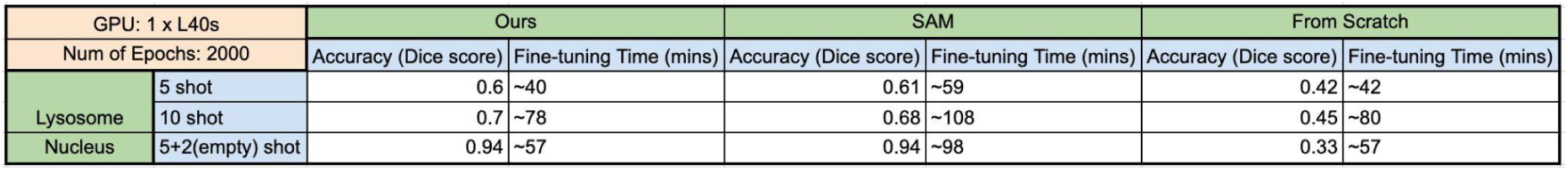
Model performance comparison. The results demonstrate that our model achieves performance comparable to SAM, with both substantially outperforming the model trained from scratch (using the same architecture as our model). The reported fine-tuning times are measured over 2000 epochs and do not reflect convergence speed. As shown in Figures 5 and 6, SAM exhibits the fastest convergence among the three models; however, its larger model size results in a longer training time per epoch and requires a larger GPU space.

Several directions remain for future investigation. First, optimizing labeling strategies is crucial, as shown in our nucleus segmentation experiments, where different strategies led to significant variations in validation Dice scores. Second, parameter-saving decisions in practical few-shot settings require careful consideration, unlike our experiments, where all slices were labeled for validation, this is not feasible in real-world scenarios. We recommend using a small maximum learning rate and gradient clipping to ensure training stability.

Additionally, training the model for 1500–2000 epochs tends to yield optimal validation performance. Overall, while SXTractor-based few-shot tomogram segmentation shows promising results, thoughtful tuning of training procedures remains key to maximizing its effectiveness.

## Methods

### 1. DINO framework

The training of SXTractor follows the original DINO [17] framework. DINO is a self-supervised knowledge distillation method that trains the student network to match the output of the teacher network (Fig. 1a) without any labels. The teacher (g_θt_) and student (g_θs_) networks have the same architecture, but with different parameters (θ_t_ and θ_s_). The teacher network is derived from the student network using an exponential moving average (EMA) of the student’s weights during training. Specifically, at each training iteration, the teacher parameters are updated as a slow-moving average of the student’s parameters:

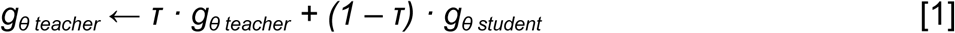

where τ is a momentum coefficient. This approach ensures the teacher provides a stable target representation over time.

During training, different augmented views (global X_global_ vs. local X_local_ views) of the same input image are passed through the student and teacher networks (global views only for the teacher network). The cropping scale of the global views is in the range of (0.4-1), and the scale for local views is (0.05-0.4). The student learns to match the teacher’s output (P_student_ and P_teacher_) representations for these views by minimizing a cross-entropy loss between their normalized outputs as [2].

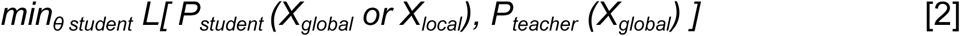

where L stands for the cross-entropy loss function. The teacher’s gradients are stopped (i.e., no backpropagation), and only the student is updated through standard gradient descent.

Over time, the student improves, and the teacher gradually incorporates this learning via the EMA update, resulting in a powerful self-supervised representation.

In the DINO framework, the standard Vision Transformer (ViT) architecture [27] is used, which will be discussed in detail in the following section. A key advantage of ViT is its ability to produce interpretable attention maps, enabling visualization of the semantic regions the model attends to during training. The spatial resolution of these attention maps is determined by the patch size used to tokenize the input image. The default patch size in DINO is 16×16, resulting in attention maps of size 64×64 for 1024×1024 input images. However, such aggressive downsampling may obscure fine-grained cellular structures. To preserve spatial details critical for identifying small organelles such as lysosomes and lipid droplets, we reduce the patch size to 8×8, thereby increasing the attention map resolution to 128×128.

### 2. Segmentation Model Architecture: SXTractor + CNN Decoder

SXTractor employs the standard Vision Transformer (ViT) architecture (ViT-S/8, ∼21 million parameters) [27]. Each input image is first divided by a manually selected patch size into non-overlapping patches, which are then flattened and linearly projected into patch embeddings. These embeddings are combined with learnable positional embeddings and passed through a series of transformer encoder layers to produce the final representation (Fig. 3, orange layers).

Different from conventional convolutional neural networks (CNNs), the Vision Transformer (ViT) processes an image as a sequence of patch tokens and produces an output of shape (B, N, C) where B stands for batch size, N is the number of patches and C is the embedding dimension - functionally analogous to feature channels in CNNs in our model. Because dimension N contains spatial information, we first permute the dimension into (B, C, N) and then reshape the N into width and length dimensions (B, C, W, H). With these modifications of the output dimensions of the vision transformer (SXTractor), a CNN-based decoder can be applied to this 2D tensor and upsample the resolution back to the input shapes (Fig. 3, green layers). The decoder consists of 3 transposed convolutional layers with ReLU as its activation function.

### 3. Segment Anything (SAM)

Segment Anything (SAM) (based on the smaller architecture ViT-B in this study, ∼93 million parameters) [10] is a foundation segmentation model trained on millions of natural images and masks. Unlike conventional architectures such as U-Net [46][47][48], SAM introduces an additional prompt encoder module alongside its image encoder and mask decoder. This design enables interactive segmentation based on user-provided prompts, such as points, bounding boxes, or masks.

However, in the context of few-shot segmentation, where no explicit prompts are provided during inference, the prompt encoder becomes functionally redundant. Therefore, we adapt SAM for few-shot learning by ignoring the prompt encoder, i.e., we do not input any form of prompts. Instead, we fine-tune the entire model end-to-end using only input images and their corresponding segmentation masks. The results indicate that pre-trained representation capabilities of SAM still provide excellent few-shot segmentation performance without any prompting.

### 4. Few-Shot Segmentation Fine-Tuning

In the few-shot segmentation setting, we conducted experiments using both 5-shot and 10-shot scenarios. Each tomogram originally contains 1024 2D slices, but many of these are empty and lack cellular structures. To make our evaluation more accurate, we removed the empty slices and retained only the 330 slices that contain meaningful cellular content. From each tomogram, we then selected 5 or 10 labeled slices as the training set for fine-tuning the model, while the remaining structure-containing slices were used as the validation (or testing) set.

To effectively fine-tune SXTractor for few-shot segmentation, careful hyperparameter selection is critical for ensuring model stability and performance. During our experiments, we explored different loss functions and observed that using only Dice loss for lysosome segmentation often resulted in convergence failure. This is likely due to Dice loss being highly sensitive to class imbalance and producing unstable gradients during early training, particularly when the predicted masks are nearly empty or lack sufficient overlap with the ground truth. To address this, we combined cross-entropy loss with Dice loss. Cross-entropy provides stronger, more stable pixel-wise supervision, especially during the initial learning phase, while Dice loss encourages better overlap and boundary alignment. This combination led to more reliable convergence and improved segmentation quality in few-shot settings.

To ensure stable training of the SXTractor-based segmentation model, we found it beneficial to use a relatively small maximum learning rate. We experimented with a range of values from 0.001 to 0.1. A maximum learning rate of 0.001 yielded more consistent validation Dice scores, while 0.1 led to faster initial convergence but often caused sharp drops in validation performance later in training. Training stability is particularly important in our setting, as we do not have the validation set for practical usage for few-shot segmentation. Therefore, a stable learning trajectory enables reliable selection of model parameters from the final epochs without overfitting or instability concerns.

We explored various data augmentation strategies to improve segmentation performance. Initial augmentations included Gaussian blur, Gaussian noise, and random horizontal and vertical flips, but these did not yield notable improvements in validation Dice scores.

Subsequently, we introduced random cropping with a crop-to-original ratio of 0.25, which resulted in a 5% increase in validation Dice score. However, this approach presents a practical drawback: during inference, each input slice must be divided into smaller patches, resulting in more complex inference operations and increased inference time. In contrast, training without random cropping yields lower validation Dice scores, but the predicted masks show no substantial qualitative differences. Thus, the decision to use random cropping should be based on the user’s priorities, specifically whether to prioritize segmentation accuracy or inference efficiency.

The GPU used in this study was NVIDIA L40s, and we calculated the time required for fine-tuning the model in different scenarios in Fig. 7 with 2000 epochs. Our selection of hyperparameters is listed below:

1. Batch size: 4 or 8
2. Optimizer: Stochastic Gradient Descent (SGD)
3. Maximum learning rate: 0.001 ∼ 0.01
4. Learning rate scheduler: One cycle scheduler
5. Loss function: Dice loss*1 + Cross entropy loss*1
6. Augmentations: Gaussian noise (probability: 0.5), Gaussian blur (probability: 0.5), Flips (probability: 0.5), Random crop (cropping area ratio: 0.25)
7. Evaluation metric: Dice score
8. Gradient clipping: 0.1

### 5. Data Description and Sample Preparation

The training data for SXTractor were collected using soft X-ray cryo-tomography through the laboratory-based soft X-ray microscope SXT-100, developed by SiriusXT. Tomograms were acquired by projecting soft X-rays through the samples across a tilt series spanning from -60° to +60° in 1° increments, with flat samples grown on TEM grids. Slices at each tilt angle were then aligned (e.g., via cross-correlation) to correct for potential thermal drift. The complete dataset comprises 4190 2D slices extracted from approximately 40 tomograms, capturing a diverse range of cellular structures.

The cell data used for segmentation fine-tuning and testing were HeLa Kyoto cells, which were seeded in Dulbecco’s Modified Eagle Medium (DMEM; Life Technologies, 31885023) containing 10% fetal bovine serum (FBS; Life Technologies, A5256801) on Quantifoil R2/2 gold mesh EM grids (Jena Biosciences, X-103-Au200). Before seeding, the grids were glow-discharged for 5 minutes in a 75:25 argon-to-oxygen atmosphere. Cells were plated at a density of 1 × 10^5^ and incubated overnight. DNA was stained with Hoechst 33342 (Sigma, 62249; 0.2 μg/mL) for 1 hour, while lysosomes were labeled using LysoTracker (Thermo Fisher, L7526; 1 μM) for 30 minutes. After washing with PBS, the samples were plunge-frozen using a Leica GP2 system with a blotting time of 6 seconds to promote vitrification and suppress ice crystal formation.

## Acknowledgements and Declarations

1. Acknowledgements

The authors gratefully acknowledge Katyja Thaysen and Professor Daniel Wüstner (Department of Biochemistry and Molecular Biology, University of Southern Denmark, DFI) for the provision of the yeast cells. We also like to thank Dr. Alessandro Zannotti (Centre for BioNano Interactions (CBNI), University College Dublin) for the provision of the VeroE6 cells, David Rogers (SiriusXT), and Stephen O’Connor (SiriusXT) for SXT data acquisition.

## 2. Declaration of Funding and Competing Interest

This work is part of the CLEXM (CORRELATIVE LIGHT, ELECTRON, AND X-RAY MICROSCOPY) consortium, a Marie Sklodowska-Curie Doctoral Networks Action (MSCA-DN), funded by the European Union under Horizon Europe [Grant agreement No. 101120151]. The authors declare no competing interests.

## 3. Data Availability Statement

The data referenced in this study are available from the corresponding author upon reasonable request.

## Author Contributions

● ShaoSen, Chueh: Methodology, Formal Analysis, Investigation, Software, Writing - original draft
● Madeleen C. Brink: Resources, Data Annotation, Qualitative Validation
● Jeremy C. Simpson: Supervision, Validation, Writing - review and editing
● Sergey Kapishnikov: Supervision, Validation, Writing - review and editing, Project Administration

## Notes

### Competing Interest Statement

The authors have declared no competing interest.

## References

[1] C. Jacobsen, “Soft x-ray microscopy,” Trends in Cell Biology, vol. 9, no. 2, pp. 44–47, Feb. 1999, doi: 10.1016/s0962-8924(98)01424-x.

[2] G. McDermott, M. A. Le Gros, C. G. Knoechel, M. Uchida, and C. A. Larabell, “Soft X-ray tomography and cryogenic light microscopy: the cool combination in cellular imaging,” Trends in Cell Biology, vol. 19, no. 11, pp. 587–595, Nov. 2009, doi: 10.1016/j.tcb.2009.08.005.

[3] K. Fahy, P. Sheridan, Sergey Kapishnikov, W. Fyans, Fergal O’Reilly, and T. McEnroe, “Laboratory-Based Soft X-ray Microscopy at a Core Facility,” Microscopy and Microanalysis, vol. 30, no. Supplement_1, Jul. 2024, doi: 10.1093/mam/ozae044.1052.

[4] O’Connor S, Rogers D, Kobylynska M, Geraets J, Thaysen K, Egebjerg JM, Brink MC, Herbsleb L, Salakova M, Fuchs L, Alves F, Feldmann C, Ekman A, Sheridan P, Fyans W, McEnroe T, O’Reily F, Fahy K, Fleck RA, Wüstner D, Simpson JC, Walter A, Kapishnikov S. 2024. Demonstrating Soft X-Ray Tomography in the lab for correlative cryogenic biological imaging using X-rays and light microscopy. bioRxiv. 10.1101/2024.12.23.629889.

[5] H. M. Hertz et al., “Laboratory cryo soft X-ray microscopy,” Journal of Structural Biology, vol. 177, no. 2, pp. 267–272, Feb. 2012, doi: 10.1016/j.jsb.2011.11.015.

[6] Z. Liu et al., “A survey on applications of deep learning in microscopy image analysis,” Computers in Biology and Medicine, vol. 134, p. 104523, Jul. 2021, doi: 10.1016/j.compbiomed.2021.104523.

[7] Y. M. Kassim, O. V. Glinskii, V. V. Glinsky, V. H. Huxley, and K. Palaniappan, “Deep Learning Segmentation for Epifluorescence Microscopy Images,” Microscopy and Microanalysis, vol. 23, no. S1, pp. 140–141, Jul. 2017, doi: 10.1017/s1431927617001386.

[8] A. F. Karimov et al., “Segmentation and classification of mast cells in histological images with deep learning,” AIP Conference Proceedings, vol. 2174, p. 020220, 2019, doi: 10.1063/1.5134371.

[9] R. Conrad and K. Narayan, “Instance segmentation of mitochondria in electron microscopy images with a generalist deep learning model trained on a diverse dataset,” Cell Systems, vol. 14, no. 1, pp. 58–71.e5, Jan. 2023, doi: 10.1016/j.cels.2022.12.006.

[10] A. Kirillov, et al., "Segment Anything," 2023 IEEE/CVF International Conference on Computer Vision (ICCV), Paris, France, 2023, pp. 3992–4003, doi: 10.1109/ICCV51070.2023.00371.

[11] Anwai Archit et al., “Segment Anything for Microscopy,” Nature Methods, Feb. 2025, doi: 10.1038/s41592-024-02580-4.

[12] J. Ma, Y. He, F. Li, L. Han, C. You, and B. Wang, “Segment anything in medical images,” Nature Communications, vol. 15, no. 1, Jan. 2024, doi: 10.1038/s41467-024-44824-z.

[13] L. Heckler, R. König and P. Bergmann, "Exploring the Importance of Pretrained Feature Extractors for Unsupervised Anomaly Detection and Localization," 2023 IEEE/CVF Conference on Computer Vision and Pattern Recognition Workshops (CVPRW), Vancouver, BC, Canada, 2023, pp. 2917–2926, doi: 10.1109/CVPRW59228.2023.00293.

[14] S. Gidaris, A. Bursuc, N. Komodakis, P. P. Pérez and M. Cord, "Boosting Few-Shot Visual Learning With Self-Supervision," 2019 IEEE/CVF International Conference on Computer Vision (ICCV), Seoul, Korea (South), 2019, pp. 8058–8067, doi: 10.1109/ICCV.2019.00815.

[15] Y. Zhou et al., “Efficient few-shot medical image segmentation via self-supervised variational autoencoder,” Medical Image Analysis, vol. 104, p. 103637, May 2025, doi: 10.1016/j.media.2025.103637.

[16] S.-C. Huang, A. Pareek, M. Jensen, M. P. Lungren, S. Yeung, and A. S. Chaudhari, “Self-supervised learning for medical image classification: a systematic review and implementation guidelines,” npj Digital Medicine, vol. 6, no. 1, pp. 1–16, Apr. 2023, doi: 10.1038/s41746-023-00811-0.

[17] M. Caron et al., "Emerging Properties in Self-Supervised Vision Transformers," 2021 IEEE/CVF International Conference on Computer Vision (ICCV), Montreal, QC, Canada, 2021, pp. 9630–9640, doi: 10.1109/ICCV48922.2021.00951.

[18] L. Ayzenberg, R. Giryes, and H. Greenspan, "DINOv2-Based Self-Supervised Learning for Few-Shot Medical Image Segmentation," 2024 IEEE International Symposium on Biomedical Imaging (ISBI), Athens, Greece, 2024, pp. 1–5, doi: 10.1109/ISBI56570.2024.10635439.

[19] S. Saha, O. Choi, and R. Whitaker, “Few-Shot Segmentation of Microscopy Images Using Gaussian Process,” Lecture notes in computer science, pp. 94–104, Jan. 2022, doi: 10.1007/978-3-031-16961-8_10.

[20] J. Dietlmeier, K. McGuinness, S. Rugonyi, T. Wilson, A. L. Nuttall, and N. E. O’Connor, “Few-shot hypercolumn-based mitochondria segmentation in cardiac and outer hair cells in focused ion beam-scanning electron microscopy (FIB-SEM) data,” Pattern Recognition Letters, vol. 128, pp. 521–528, Dec. 2019, doi: 10.1016/j.patrec.2019.10.031.

[21] Y. Dawoud, J. Hornauer, G. Carneiro, and Vasileios Belagiannis, “Few-Shot Microscopy Image Cell Segmentation,” Lecture notes in computer science, pp. 139–154, Jan. 2021, doi: 10.1007/978-3-030-67670-4_9.

[22] P.-Y. Tung and R. J. Harrison, “Efficient microstructure segmentation in three-dimensional imaging: Combining few-shot learning with the segment anything model,” Next Materials, vol. 8, p. 100663, Apr. 2025, doi: 10.1016/j.nxmate.2025.100663.

[23] S. Akers et al., “Rapid and flexible segmentation of electron microscopy data using few-shot machine learning,” npj Computational Materials, vol. 7, no. 1, Nov. 2021, doi: 10.1038/s41524-021-00652-z.

[24] Z. Shen, Z. Liu, J. Qin, L. Huang, K.-T. Cheng, and M. Savvides, “S2-BNN: Bridging the Gap Between Self-Supervised Real and 1-bit Neural Networks via Guided Distribution Calibration,” 2021 IEEE/CVF Conference on Computer Vision and Pattern Recognition (CVPR), pp. 2165–2174, Jun. 2021, doi: 10.1109/cvpr46437.2021.00220.

[25] C. Ting, K. Simon, S. Kevin, N. Mohammad, and H. E. Geoffrey, “Big Self-Supervised Models are Strong Semi-Supervised Learners,” Advances in Neural Information Processing Systems, vol. 33, 2020, Accessed: Nov. 20, 2022. [Online]. Available: https://proceedings.neurips.cc/paper/2020/hash/fcbc95ccdd551da181207c0c1400c655-Abstract.html

[26] M. Noroozi, A. Vinjimoor, P. Favaro and H. Pirsiavash, "Boosting Self-Supervised Learning via Knowledge Transfer," 2018 IEEE/CVF Conference on Computer Vision and Pattern Recognition, Salt Lake City, UT, USA, 2018, pp. 9359–9367, doi: 10.1109/CVPR.2018.00975.

[27] A. Dosovitskiy et al., “An Image is Worth 16x16 Words: Transformers for Image Recognition at Scale,” *arXiv.org*, Jun. 03, 2021. https://www.arxiv.org/abs/2010.11929v2

[28] J. Liu et al., “Locating the ‘missing wedge’ artifacts from limited-angle CT reconstruction,” Microscopy and Microanalysis, vol. 24, no. S2, pp. 140–141, Aug. 2018, doi: 10.1017/s1431927618013089.

[29] A. Hosna, E. Merry, J. Gyalmo, Z. Alom, Z. Aung, and M. A. Azim, “Transfer learning: a friendly introduction,” Journal of Big Data, vol. 9, no. 1, Oct. 2022, doi: 10.1186/s40537-022-00652-w.

[30] F. Li et al., "Mask DINO: Towards A Unified Transformer-based Framework for Object Detection and Segmentation," 2023 IEEE/CVF Conference on Computer Vision and Pattern Recognition (CVPR), Vancouver, BC, Canada, 2023, pp. 3041–3050, doi: 10.1109/CVPR52729.2023.00297.

[31] S. Liu et al., “Grounding DINO: Marrying DINO with Grounded Pre-training for Open-Set Object Detection,” Lecture notes in computer science, pp. 38–55, Nov. 2024, doi: 10.1007/978-3-031-72970-6_3.

[32] Y. Tian et al., "Fast-iTPN: Integrally Pre-Trained Transformer Pyramid Network With Token Migration," in IEEE Transactions on Pattern Analysis and Machine Intelligence, vol. 46, no. 12, pp. 9766–9779, Dec. 2024, doi: 10.1109/TPAMI.2024.3429508.

[33] Y. LeCun et al., “Handwritten Digit Recognition with a Back-Propagation Network,” Neural Information Processing Systems, 1989. https://proceedings.neurips.cc/paper/1989/hash/53c3bce66e43be4f209556518c2fcb54-Abstr act.html

[34] A. Krizhevsky, I. Sutskever, and G. E. Hinton, “ImageNet Classification with Deep Convolutional Neural Networks,” Communications of the ACM, vol. 60, no. 6, pp. 84–90, May 2012, Available: https://proceedings.neurips.cc/paper_files/paper/2012/file/c399862d3b9d6b76c8436e924a68c45b-Paper.pdf

[35] K. Simonyan and A. Zisserman, “Very deep convolutional networks for large-scale image recognition,” 3rd International Conference on Learning Representations (ICLR 2015), 2015, Available: https://ora.ox.ac.uk/objects/uuid:60713f18-a6d1-4d97-8f45-b60ad8aebbce

[36] C. Szegedy et al., "Going deeper with convolutions," 2015 IEEE Conference on Computer Vision and Pattern Recognition (CVPR), Boston, MA, USA, 2015, pp. 1–9, doi: 10.1109/CVPR.2015.7298594.

[37] K. He, X. Zhang, S. Ren and J. Sun, "Deep Residual Learning for Image Recognition," 2016 IEEE Conference on Computer Vision and Pattern Recognition (CVPR), Las Vegas, NV, USA, 2016, pp. 770–778, doi: 10.1109/CVPR.2016.90.

[38] M. Abu et al., “A Comprehensive Performance Analysis of Transfer Learning Optimization in Visual Field Defect Classification,” Diagnostics, vol. 12, no. 5, p. 1258, May 2022, doi: 10.3390/diagnostics12051258.

[39] A. Newell and J. Deng, "How Useful Is Self-Supervised Pretraining for Visual Tasks?," 2020 IEEE/CVF Conference on Computer Vision and Pattern Recognition (CVPR), Seattle, WA, USA, 2020, pp. 7343–7352, doi: 10.1109/CVPR42600.2020.00737.

[40] J. Guo and C. A. Larabell, “Soft X-ray tomography: virtual sculptures from cell cultures,” Current Opinion in Structural Biology, vol. 58, pp. 324–332, Oct. 2019, doi: 10.1016/j.sbi.2019.06.012.

[41] M. Harkiolaki, M. C. Darrow, M. C. Spink, E. Kosior, K. Dent, and E. Duke, “Cryo-soft X-ray tomography: using soft X-rays to explore the ultrastructure of whole cells,” Emerging Topics in Life Sciences, vol. 2, no. 1, pp. 81–92, Mar. 2018, doi: 10.1042/etls20170086.

[42] J.-H. Chen et al., “A protocol for full-rotation soft X-ray tomography of single cells,” STAR Protocols, vol. 3, no. 1, p. 101176, Feb. 2022, doi: 10.1016/j.xpro.2022.101176.

[43] V. Loconte et al., “Using soft X-ray tomography for rapid whole-cell quantitative imaging of SARS-CoV-2-infected cells,” Cell Reports Methods, vol. 1, no. 7, p. 100117, Nov. 2021, doi: 10.1016/j.crmeth.2021.100117.

[44] A. Erozan, P. D. Lösel, V. Heuveline, and V. Weinhardt, “Automated 3D cytoplasm segmentation in soft X-ray tomography,” iScience, vol. 27, no. 6, p. 109856, Jun. 2024, doi: 10.1016/j.isci.2024.109856.

[45] Ayse Erozan, A. Zeilmann, Anthoula Chatzimpinou, Fariha Mahzabin Annesha, V. Heuveline, and V. Weinhardt, “Spatially-Informed Multi-Stage Active Learning for Segmentation of Nucleus in Soft X-ray Tomography,” pp. 1–7, Mar. 2025, doi: 10.1109/cism64958.2025.11060865.

[46] O. Ronneberger, P. Fischer, and T. Brox, “U-Net: Convolutional Networks for Biomedical Image Segmentation,” Lecture Notes in Computer Science, vol. 9351, pp. 234–241, 2015, doi: 10.1007/978-3-319-24574-4_28.

[47] F. Isensee, P. F. Jaeger, S. A. A. Kohl, J. Petersen, and K. H. Maier-Hein, “nnU-Net: a self-configuring method for deep learning-based biomedical image segmentation,” Nature Methods, vol. 18, no. 2, pp. 203–211, Dec. 2020, doi: 10.1038/s41592-020-01008-z.

[48] Ö. Çiçek, A. Abdulkadir, S. S. Lienkamp, T. Brox, and O. Ronneberger, “3D U-Net: Learning Dense Volumetric Segmentation from Sparse Annotation,” Medical Image Computing and Computer-Assisted Intervention – MICCAI 2016, vol. 9901, pp. 424–432, 2016, doi: 10.1007/978-3-319-46723-8_49.

